# Bacterial persisters in long-term infection: emergence and fitness in a complex host environment

**DOI:** 10.1101/561589

**Authors:** Jennifer A. Bartell, David R. Cameron, Biljana Mojsoska, Janus Anders Juul Haagensen, Lea M. Sommer, Kim Lewis, Søren Molin, Helle Krogh Johansen

**Author notes:** Department of Science and Environment, Roskilde University, Roskilde, Denmark (Biljana Mojsoska). Contributed equally to the work and listed alphabetically (Jennifer A. Bartell, David R. Cameron, Biljana Mojsoska).

## Abstract

Despite intensive antibiotic treatment, *Pseudomonas aeruginosa* often persists in the airways of cystic fibrosis (CF) patients for decades, and can do so without antibiotic resistance development. Using high-throughput screening assays of bacterial survival after treatment with high concentrations of ciprofloxacin and tobramycin, we have determined the prevalence of persisters in a large patient cohort consisting of 460 longitudinal isolates of *P. aeruginosa* from 39 CF patients. Thirty patients exhibited high persister variants (Hip, defined by survival of at least 75% of replicates) in at least one of the two antibiotic screens (25% of isolates in total). Few bacterial lineages were dominated by Hip, but Hip emergence increased over lineage colonization time. Furthermore, transient lineages were significantly less likely to exhibit Hips than non-transient lineages, suggesting that the Hip phenotype is decisive for long-term establishment of a lineage. While we observed no strong signal of adaptive genetic convergence across all lineages with Hip emergence, Hip+ lineages were significantly correlated with lineages with slow growing isolates. Finally, we evaluated Hips in a model CF structured environment by testing the fitness properties of otherwise genotypically and phenotypically similar low-persister (Lop) and Hip isolates in co-culture using a flow-cell biofilm system with antibiotic dosing modelled on *in vivo* dynamics. Hip survived ciprofloxacin treatment better than Lop. Our results strongly argue against the existence of a single dominant molecular mechanism underlying bacterial antibiotic persistence. We instead show that many routes, both phenotypic and genetic, are available for persister formation and consequent increases in strain fitness in CF airways.

**BIOLOGICAL SCIENCES:** Microbiology

## Introduction

Antibiotic-tolerant persister cells are suspected to be a significant clinical problem that has been seriously neglected in favor of combating antibiotic-resistant bacteria, though persisters were in fact described shortly after the clinical introduction of antibiotics (1). Persisters are distinct from antibiotic-resistant mutants, as they do not grow in the presence of antibiotics. Instead, they remain dormant during antibiotic exposure but retain the capacity to resuscitate and restore the population when antibiotic concentrations drop (2–4). However, our understanding of the physiology and clinical relevance of persister cells is limited, given the difficulty in reliably isolating what is theorized to be a stochastic phenotype *in vitro*, much less monitoring this phenotype in routine clinical care. Thus, while a few characterizations of small environmental isolate collections have shown that formation of persisters varies across strains (5, 6), few studies have assayed persister formation in clinical or other complex environmental scenarios. One study of oral carriage (0-19 weeks) of *Candida albicans* isolates from 22 cancer patients undergoing chemotherapy found that patients with carriage of greater than 8 weeks had significantly higher persister levels than those with less than 8 weeks of carriage, but did not address the underlying mechanisms of persistence in this pathogen (7). To examine the underpinnings and long-term impact of the persister phenotype in a clinical scenario, both a large, aligned patient cohort that places the bacteria under similar environmental stresses as well as isolate sampling at a resolution that captures the emergence and longevity of the phenotype are needed.

*P. aeruginosa* is the most frequent cause of chronic airway infections in patients with CF (8, 9). Mutations in the cystic fibrosis transmembrane conductance regulator (*CFTR*) gene often result in inefficient mucociliary clearance of bacteria from the airways, creating opportunities for bacterial colonization (10, 11). Upon entering the host, environmental *P. aeruginosa* adapts to the CF lung environment, ultimately establishing an incurable airway infection (12, 13). Despite intensive antibiotic treatment from the first discovery of the bacterium in the lung, resistance emergence in the first years of infection is surprisingly low (14, 15). In the absence of clinically defined antibiotic resistance, survival of the bacteria is likely enabled by diverse and co-occurring traits including slowed growth rate, biofilm formation, and the production of small fractions of antibiotic tolerant subpopulations (16–18). How persister cells interrelate with these complex and co-selected changes *in vivo* is rarely accounted for in *in vitro* persister studies, but is likely clinically important.

These factors also complicate the search for genetic mechanisms of the persister phenotype that are clinically impactful. While persister cells are stochastic phenotypic variants in any bacterial population, genetic changes in bacterial populations have been shown to produce a high persister state, producing increased numbers of antibiotic tolerant cells following exposure to antibiotics in *in vitro* studies of pathogenic species (19, 20). Some of these genetic changes have also been observed in clinical isolates; within a set of 477 commensal or urinary tract infection isolates of *Escherichia coli*, 24 exhibited a mutation in the canonical persister gene *hipA*, and the causality between a *hipA7* mutation and a Hip phenotype confirmed by deleting this allele from one of the clinical isolates (21). An investigation in young CF patients showed an increase in persister phenotype in early/late infection isolate pairs from 14 patients. In this study, 35 longitudinal *P. aeruginosa* isolates taken from one child over a 96-month period showed increased levels of persister cells over time as well as an accumulation of 68 mutations between the first and last isolate (16). However, the mutations in the single patient resembled those known to accumulate in other CF patients over infection rather than any mutations previously associated with the Hip phenotype in persister-focused *in vitro* studies.

To acquire a high-resolution pan-cohort perspective of persister emergence, genetic mechanism, and impact in long-term infections, we have screened 460 longitudinal isolates of *P. aeruginosa* collected from 39 young CF patients over a 10-year period from early colonization onward for high persister variants (Hip, defined by survival of at least 75% of replicates) tolerant to two unrelated antibiotics (ciprofloxacin and tobramycin). This unique isolate collection allows us to determine Hip prevalence and dynamics during each colonizing strain’s transition from environmental isolate to persistent pathogen. We describe relationships between the Hip phenotype and response to each drug, the age of the isolate, and other adaptive traits in longitudinal infections. We show that the Hip phenotype, defined in this study as a strong and reliable recovery from antibiotic challenge that is a serious concern for the clinic, is an independent and widespread trait. We further search for genetic and phenotypic changes associated with the Hip phenotype in independent clonal lineages within distinct patients, which may suggest adaptive routes to producing this phenotype. Finally, we show that the Hip phenotype generally accumulates over time in patients via several archetypal patterns, appears to contribute to long-term persistence of lineages, and increases the fitness of colonizing populations of *P. aeruginosa* in antibiotic-treated CF patient lungs.

## Results

### The isolate collection

We examined a collection of 460 *P. aeruginosa* airway isolates obtained from 39 young CF patients over a 10 year period while they were treated at the Copenhagen CF Centre at Rigshospitalet (22). These patients represent a cohort aligned at the early infection stage and undergoing similar treatment regimens per CF Centre guidelines, with repeated culture of *P. aeruginosa* from their monthly sputum sampling within a time frame of 2-10 years (patient inclusion was on a rolling basis over the study period in order to capture all early colonization cases). Early isolates therefore represent bacteria that have not been exposed to substantial antibiotic treatment before the study start excepting rare cases of strain transmission from another patient. The bacterial CF isolates have been grouped into 52 genetically distinct clone types (22), and while many patients retained a monoclonal infection during the entire course of infection, half (n=20, 51.3 %) were infected at least transiently with another clone type. To effectively account for these multi-clonal infections, clinical isolates are described by their patient-specific lineage combining the clone type and the patient of origin (74 lineages in total). Throughout this paper, we will also refer to ‘Time since first detection’ for each isolate, which represents the length of time between first detection and subsequent isolations of the same patient-specific lineage.

### Identification of high persister (Hip) isolates by high-throughput screening

We screened the collection of *P. aeruginosa* isolates for the propensity to survive in the presence of high concentrations of antibiotics. We chose two distinct antibiotics that matched the following criteria; (i) they are frequently used to treat early *P. aeruginosa* infections in CF patients; (ii) have contrasting targets and mechanisms of action; (iii) drive resistance development with different dynamics in patients; and (iv) are each bactericidal toward stationary phase *P. aeruginosa* (23, 24). On this basis, we performed two independent screens using either the fluoroquinolone ciprofloxacin, which interacts with DNA gyrase, or the aminoglycoside tobramycin, which acts upon bacterial ribosomes. Briefly, *P. aeruginosa* subcultures in micro-titer plates were grown for 48 hours until they reached stationary phase, after which they were challenged with antibiotics (100 μg/ml) for 24 hours before survival was assessed (Fig. 1A). A standardised antibiotic concentration was used that was at least 25-times higher than the European Committee on Antimicrobial Susceptibility Testing (EUCAST) resistance breakpoint for each drug, minimising the chance that the screen selected for isolates with modestly elevated minimum inhibitory concentrations (MICs). Within each antibiotic screen, isolates were assayed eight times (technical quadruplicates performed in duplicate biological experiments with a positive growth control for at least 3 of 4 replicates in each experiment) and scored based on the capacity to re-grow after antibiotic treatment. An isolate was given a score of 0 if it failed to re-grow in any replicate of an experiment, a score of 1 if it grew once in both biological duplicates, a score of 2 if it grew in half of the technical replicates in each experiment, a score of 3 if it grew in at least three replicates in each experiment, and a score of 4 if it grew in all replicates (Fig. 1B, scores for the ciprofloxacin screen). To validate our high-throughput screening approach, we selected 25 isolates, and enumerated CFUs following 24 hours of ciprofloxacin treatment using standard survival assays. We observed a significant positive correlation between persister score and CFU/ml following treatment (r_2_ 0.5719, p < 0.0001, SI Appendix, Fig. S1), thus demonstrating the validity of our experimental approach.

**Figure 1.**
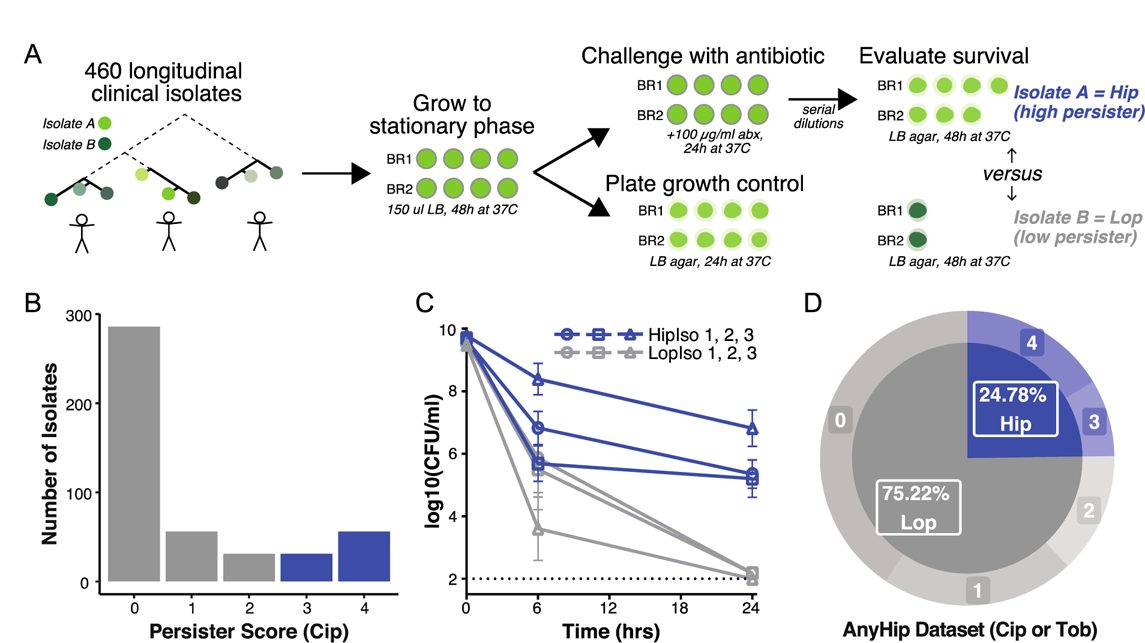
High-throughput screening approach for isolates with a high persister (Hip) phenotype. (A) A large collection of *Pseudomonas aeruginosa* clinical isolates were grown to stationary phase in quadruplicate wells for two biological replicate (BR) experiments (each isolate tested 8 times in total). The screen was performed twice, using either ciprofloxacin or tobramycin. Each isolate was treated with 100 μg/ml of antibiotic for 24 hours, while growth was assessed by plating on LB agar. Following antibiotic treatment, cultures were diluted then plated on agar, at which point survival was assessed. Each isolate was given a persister score based on consistent replicate survival following treatment. Isolates for which 3-4 replicates survived for each BR were given a score of 3-4, respectively, and were considered high persisters (Hip). Isolates with a respective score of 0-2 were considered low persisters (Lop). **(B)** Score distribution of *P. aeruginosa* Hip (blue) and Lop (grey) isolates against ciprofloxacin. **(C)** Traditional time-kill assays were performed for three Hip (HipIso 1, 2, 3) and three Lop isolates (LopIso 1, 2, 3) from the same patient to validate the high throughput screen. Colony forming units (CFU) per ml were determined following treatment with 100 μg/ml of ciprofloxacin. Data are the mean of 6 independent cultures, bars represent SEM. **(D)** Distribution of *P. aeruginosa* Hip (blue) and Lop (grey) isolates for the AnyHip dataset, where Hips were given a score of 3 or 4 in at least one of the antibiotic screens.

We defined high persister (Hip) isolates as those scoring either 3 or 4 (i.e. at least six of eight technical replicates re-grew) and low persister (Lop) isolates as those scoring between 0 and 2. This stringent scoring system was used to minimise the mis-classification of false Hips and focus our analysis on isolates reliably producing persister cells, representing the most concerning phenotype in a clinical environment. Isolates with a score of 4 made up the largest Hip group, while the largest Lop group consisted of isolates scored as 0, failing to grow in any replicate (Fig 1B). To validate this classification system, we selected six isolates from the same patients, three of which were putative Hips, and three of which were classed as Lops and performed time-dependent killing assays. The isolates displayed typical, biphasic killing (Fig. 1C). Each of the three putative Hip strains harboured a greater subpopulation of surviving persister cells, thus they represent true Hips as defined by recent guidelines (25). In total, 24.8% of screened isolates (114 isolates) exhibited a persister phenotype in at least one of the screens (Fig. 1D, AnyHip dataset). In Figure 2, we show the distribution of Hips across our isolates with respect to (A) ciprofloxacin persistence (87 Cip Hip isolates), (B) tobramycin persistence (60 Tob Hip isolates), and (C) multi-drug persistence (33 MD Hip isolates), where an isolate scored 3 or more in both screens (Dataset 1). The bottom panel of Figure 2C shows an explicit breakdown of dataset overlap.

**Figure 2.**
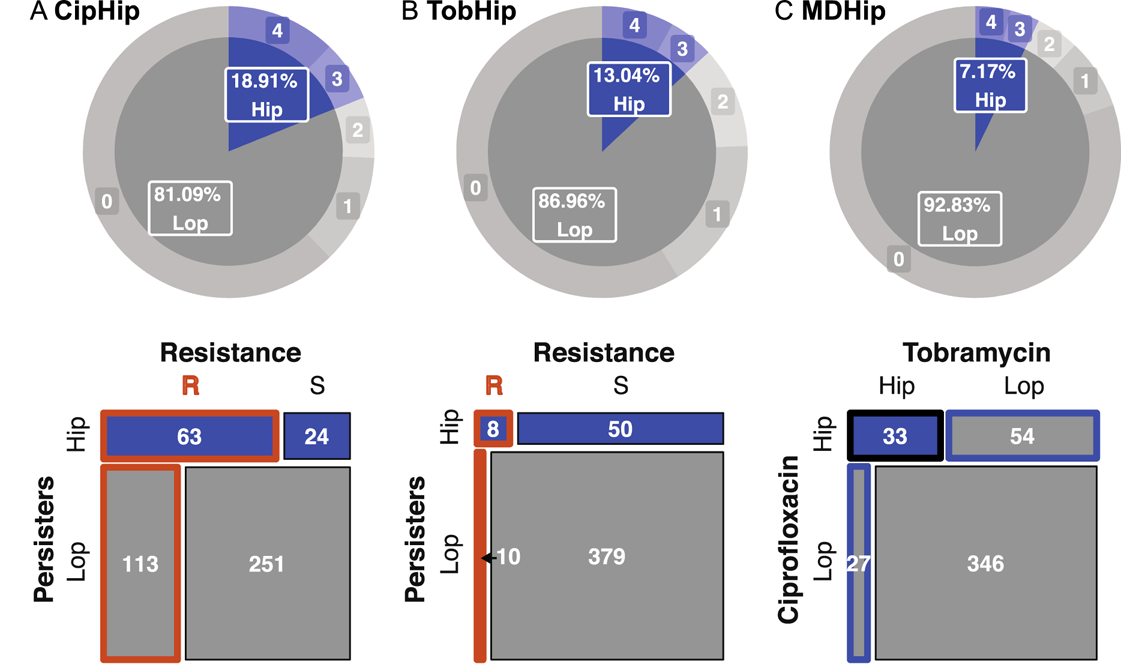
Persister screening results. **(A)** Distribution of Hip (blue) and Lop (grey) isolates following ciprofloxacin (CipHip,) or **(B)** tobramycin treatment (TobHip), and **(C)** the overlap between these datasets (MDHip - multi-drug Hip). For **A** and **B**, the bottom panel shows a mosaic plot (multi-way contingency table) for the comparison of isolate persister class versus resistance class for each antibiotic. Hip and Lop persister isolates were classified as susceptible (S) or resistant (R) according to EUCAST breakpoints (cip: S≤0.5µg/ml, tob: S≤4µg/ml) based on their MIC obtained via E-test. The area of each cell is proportional to the frequency of isolates with the indicated combination of Hip and resistance classification, and resistance-associated cells are further highlighted by orange borders. The bottom panel of **C** shows the overlap of Hip and Lop isolates under ciprofloxacin versus tobramycin treatment. MDHip are highlighted by a black border.

### The persister phenotype is antibiotic-specific and is dissociated from antibiotic resistance

As we were using antibiotics used in patient treatment to select for persister isolates, we evaluated if we were simply selecting for resistant isolates as opposed to true Hips. Ciprofloxacin and tobramycin MICs were determined using E-tests for isolates within the collection. Most of the isolates were characterized as susceptible based on the EUCAST breakpoints (Fig. 2A, B). Twenty-four ciprofloxacin susceptible isolates were Hip while 113 Lop variants were resistant. Similarly, 10 tobramycin resistant isolates were Lop while 50 Hip isolates were tobramycin susceptible. We then contrasted the emergence of antibiotic resistance versus Hip for all study lineages (Table S1). Of the 74 lineages assessed, resistance emerged without Hip detection in 14 lineages (40%), while Hips emerged without resistance in 16 lineages. Resistance and persistence emerged simultaneously in 11 lineages. Resistance preceded Hip in 9 lineages, while Hip preceded resistance in 5 lineages. If we used a less stringent persister classification (persister score greater than 0 instead of 2), we observed that resistance precedes persistence in 5 lineages, emerges simultaneously in 7 lineages, and follows persistence in 15 lineages. Taken together, these data confirm that our screening approach identified a persister phenotype separate from an antibiotic-resistant phenotype.

### The persister phenotype is enriched in patient-specific lineages with slow-growing isolates

Along with changes in antibiotic susceptibility, the persister phenotype in our collection is not arising in isolation from other adaptations. We and others have previously observed that CF isolates adapt towards slow growth rates, increased resistance to antibiotics, and preference for a biofilm lifestyle (18, 26, 27). A specific association between slowing growth rate and the Hip phenotype has also been proposed (28). To probe interrelationships with other phenotypes, we used a principle component analysis to evaluate the distribution of Hip (blue diamonds) versus Lop (grey circles) variants by multiple traits under selection pressure in the CF lung. We see that Any Hip variants group with isolates exhibiting more adapted traits (increased antibiotic MICs and slowing growth), but they also appear across the full phenotypic space alongside Lop isolates (Fig. 3A).

**Figure 3.**
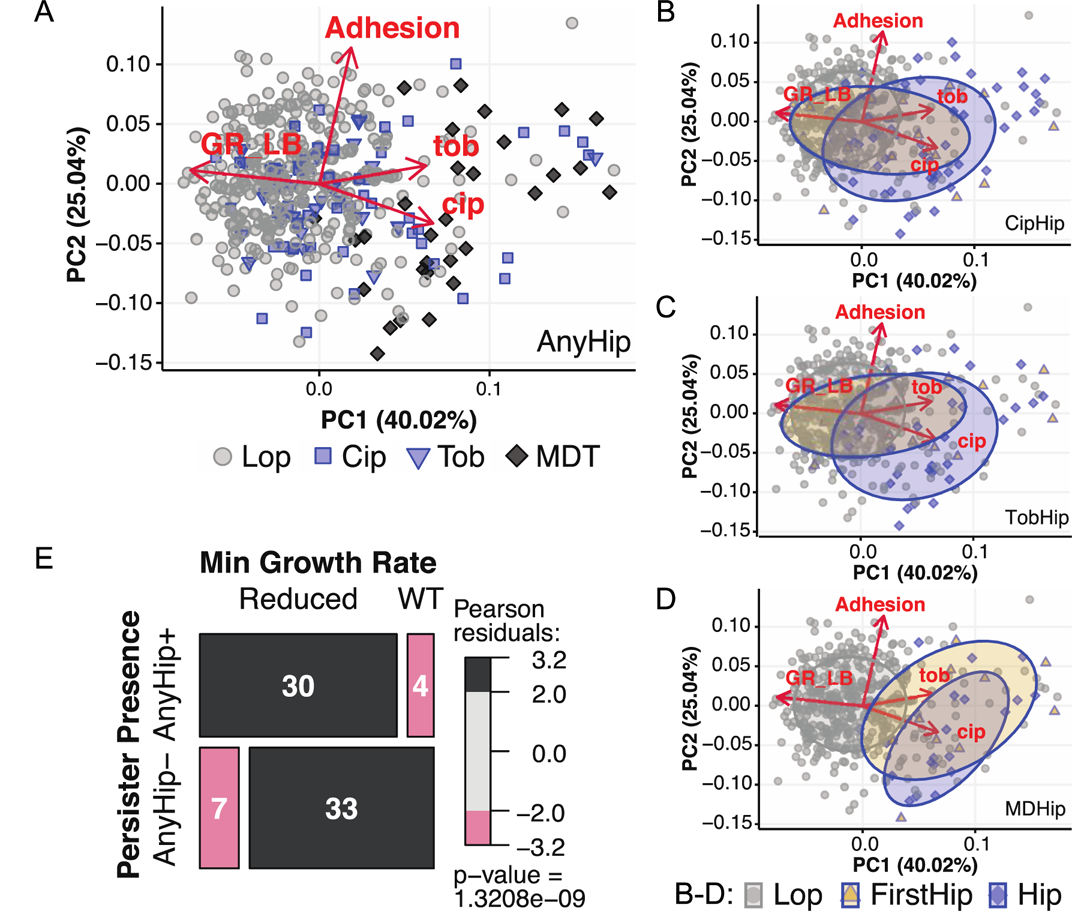
High persisters in the multi-trait landscape. **(A)** Lop (grey circle), Cip Hip (blue squares), Tob Hip (blue triangles) and MD Hip isolates (black diamonds) were analyzed via principle component analysis with respect to their similarity with other infection-linked traits: growth rate (GR_LB), adhesion, ciprofloxacin MIC (cip) and tobramycin MIC (tob). 446 isolates with complete trait sets were included. Hip isolates do not consistently cluster with any one additional trait. For **(B)** Cip Hip, **(C)** Tob Hip and **(D)** MDHip, the first Hip isolates from a lineage (FirstHip, yellow triangles) were highlighted as Hip variants with mitigated effects of other accumulating mutations within the lineage to improve cross-lineage comparison. In each case, FirstHip and the remaining Hip isolates shift to various degrees from ‘naïve’ towards ‘adapted’ levels given the particular Hip dataset. We illustrate this using data ellipse enclosing samples approximately within the first standard deviation (t distribution, 68% of the set) for isolate sets characterized as FirstHip (yellow ellipse), and the remaining Hips (blue ellipse). **(E)** We visualized the association between lineages that produced Hips versus slow growing isolates (identified by the minimum growth rate of lineage isolates falling below 75% of the *P. aeruginosa* PAO1 growth rate based on a 45 minute generation time in LB). Association between variables is illustrated by a mosaic plot (multi-way contingency table visualization) where color indicates significant deviation from the expected frequency of lineages in each cell under trait independence using Pearson’s chi-squared test.

When comparing the ciprofloxacin versus tobramycin screen data (Fig. 3B-C), the data ellipses enclosing the approximated majority of each population (68% of the population, t distribution) show that Hip variants (blue ellipse) only partially separate from Lop variants (grey ellipse). If we assess the first Hip variant of each lineage (FirstHip – blue ellipse with yellow fill), we see FirstHips overlap substantially with both Lops and Hips. This variation of initial adaptive state could be due to different adaptive trajectories with patients as well as lapses of time between

Hip emergence and isolation. The MD FirstHips show a more distinctive localization away from Lops, which suggests further benefits of serious growth defects but could also be the result of increased colonization time necessary to evolve both persister traits (Fig. 3D). If we look at the likelihood of each lineage containing both Hips and slow growing isolates (where the minimum growth rate of the lineage is less than 75% of the growth rate of PAO1), we see a significant relationship between incidence of slow growing isolates and AnyHip isolates (Fig. 3E). In total, these results emphasize the complexity of selection pressures at play, resulting in concurrent adaptation of distinct traits that may both influence each other and have related genetic underpinnings.

### Evolution of the persister phenotype is not genetically convergent across patient-specific lineages

Sequencing and identification of genetic variations accumulating within each clone type have been previously performed for most of the isolates used in this study (403 isolates, 46 lineages). Genes targeted in convergent evolution were identified by the significant enrichment of observed lineages with mutations in those genes compared to the number of lineages expected to have mutations in the same genes according to genetic drift (derived from a simulated evolution where lineages accumulate an equivalent number of mutations randomly for 1000 independent evolution simulations) (22). We split our dataset into Hip and Lop variants, and then performed this same observed versus expected lineage enrichment analysis for each population (see Materials and Methods for further details). The analysis was performed using each of four datasets (ciprofloxacin Hip, tobramycin Hip, MD Hip and Any Hip), and the ratio of lineage enrichment of mutated genes for Hip versus Lop variants allowed us to identify candidate ‘*hip*’ genes for each set (Table 1 and Dataset 2). For completeness, we also performed an additional genetic analysis focusing on mutations in non-coding sequences (Dataset 2).

**Table 1.**
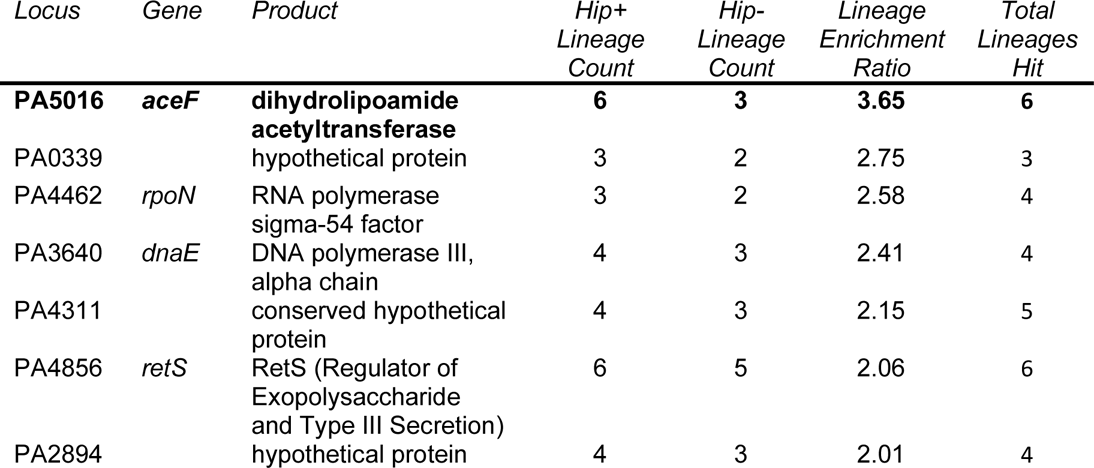
Lineage-based mutation enrichment analysis. Mutated genes enriched in AnyHip versus Lop dataset as assessed from a convergent evolution perspective accounting for lineage adaptation. Lineage enrichment ratio was calculated by dividing lineage-based gene mutation enrichment within Hip variants by that within Lop variants for each gene. Top Hip-linked genes were selected via the following criteria: greater than 2 lineages presenting mutations in that gene in the Hip population and a lineage enrichment ratio greater than 2. The only gene also enriched in MDHip lineages, *aceF*, is in boldface.

| Locus | Gene | Product | Hip+<br>Lineage<br>Count | Hip-<br>Lineage<br>Count | Lineage<br>Enrichment<br>Ratio | Total<br>Lineages<br>Hit |
| --- | --- | --- | --- | --- | --- | --- |
| **PA5016** | ***aceF*** | **dihydrolipoamide**<br>**acetyltransferase** | **6** | **3** | **3.65** | **6** |
| PA0339 |  | hypothetical protein | 3 | 2 | 2.75 | 3 |
| PA4462 | *rpoN* | RNA polymerase<br>sigma-54 factor | 3 | 2 | 2.58 | 4 |
| PA3640 | *dnaE* | DNA polymerase III,<br>alpha chain | 4 | 3 | 2.41 | 4 |
| PA4311 |  | conserved hypothetical<br>protein | 4 | 3 | 2.15 | 5 |
| PA4856 | *retS* | RetS (Regulator of<br>Exopolysaccharide<br>and Type III Secretion) | 6 | 5 | 2.06 | 6 |
| PA2894 |  | hypothetical protein | 4 | 3 | 2.01 | 4 |

In general, searching for *hip* genes accumulating non-synonymous mutations in Hip+ lineages revealed only a weak signal for convergent evolution. Only one lineage assessed in the genetic screen had only Hips present (the only isolate of the lineage assessed in our screen), so practically all lineages with Hip isolates (Hip+ lineages, 29 included in the genetic study) also contained Lops. Thus, mutated genes that were enriched 2-3 fold in independently evolved Hip+ lineages were also frequently present in Lop isolates of the same lineage. Our lineage enrichment ratio ultimately identified 12 mutated genes enriched in ciprofloxacin Hip+ lineages (SI Appendix, Dataset 2), 13 mutated genes in tobramycin Hip+ lineages (SI Appendix, Dataset 2), a solitary gene for MD Hip+ lineages (SI Appendix, Dataset 2), and 7 mutated genes for Any Hip+ lineages (Table 1).

Of note, there was a surprising lack of the most prominent ‘*hip*’ genes previously identified in *in vitro* studies and screens of *P. aeruginosa* (SI Appendix, Table S2). None of the lineage enrichment data pointed toward RNA endonuclease-type toxin-antitoxin systems under adaptive selection, which supports recent research that has questioned the contribution of these systems to persistence in numerous bacterial pathogens (29–31). Instead, they belonged to diverse functional categories including transcriptional regulation/two component regulatory systems (4 genes), energy metabolism (3 genes), and DNA replication and repair (2 genes). In the Any Hip analysis as well as the MD Hip analysis, the shared top gene target, *aceF,* encodes a component of the pyruvate dehydrogenase complex, a key player in central metabolism, for which functional mutations are known to reduce growth rate (32–34). Other genes in Table 1 include major regulators *rpoN*, known to induce a growth defect when functionally mutated (35), and *retS*, which when functionally mutated induces an array of phenotypic changes linked to chronic infection such as a non-motile biofilm lifestyle (36, 37). These genes, as well as hypothetical protein PA4311, also overlap with the ‘pathoadaptive’ mutationally enriched gene list identified in our prior study of convergent evolution across all lineages (22).

### Hip variants emerge via diverse incidence patterns

The lack of strong genetic signatures differentiating Hip from Lop isolates motivated us to examine the temporal dynamics of high persister incidence using our comprehensive Any Hip dataset. In half of the patients, the earliest bacterial isolate is also the first-ever identified *P. aeruginosa* in the clinic and the other patients’ isolates also cover most of the initial colonization phase. We can thus estimate the emergence of the Hip phenotype as *P. aeruginosa* adapts from a wild type-similar naïve state into an adapted persistent pathogen. Previous findings have indicated that the number of Hip variants from a lineage may increase over time as the bacteria adapt to the antibiotic pressure in the host, and that once a Hip isolate is observed, it is assumed to persist in the infecting population of the patient (7, 16).

To illustrate the range of persister dynamics we observe, we grouped each lineage by an array of descriptors. The lineage descriptors include Hip presence versus absence (Hip+ vs Hip-), transience of the lineage (whether it appears for less than 2 years, less than half the length of a patient’s infection and is afterwards replaced by another lineage), continuity of Hip variants (whether Hips are present for at least 3 sampling dates in a row), dominance of Hip variants (whether Hips make up at least 2/3 of all isolates of a lineage), and whether a Hip variant initiates the lineage. Figure 4A shows the ordered distribution of the lineages in 10 different groups based on descriptor sets, illustrating the diversity of lineage Hip dynamics. We see that: 1) 34 of 74 lineages are Hip+, 2) 28 of 40 Hip-lineages are transient, while 4 Hip+ lineages are transient, 3) 21 Hip+ lineages have Hip variants appear irregularly versus 7 lineages that exhibit continuous periods and/or dominance of Hip variants, and 4) 12 lineages have initiating Hip+ variants. Thus, the fraction (24.8%, Fig. 1D) of total isolates with a Hip phenotype appears to be distributed over a subset of lineages (45.9%) in both stable (continuous/dominant) and stochastic patterns of incidence, rather than present in every evolving lineage.

**Figure 4.**
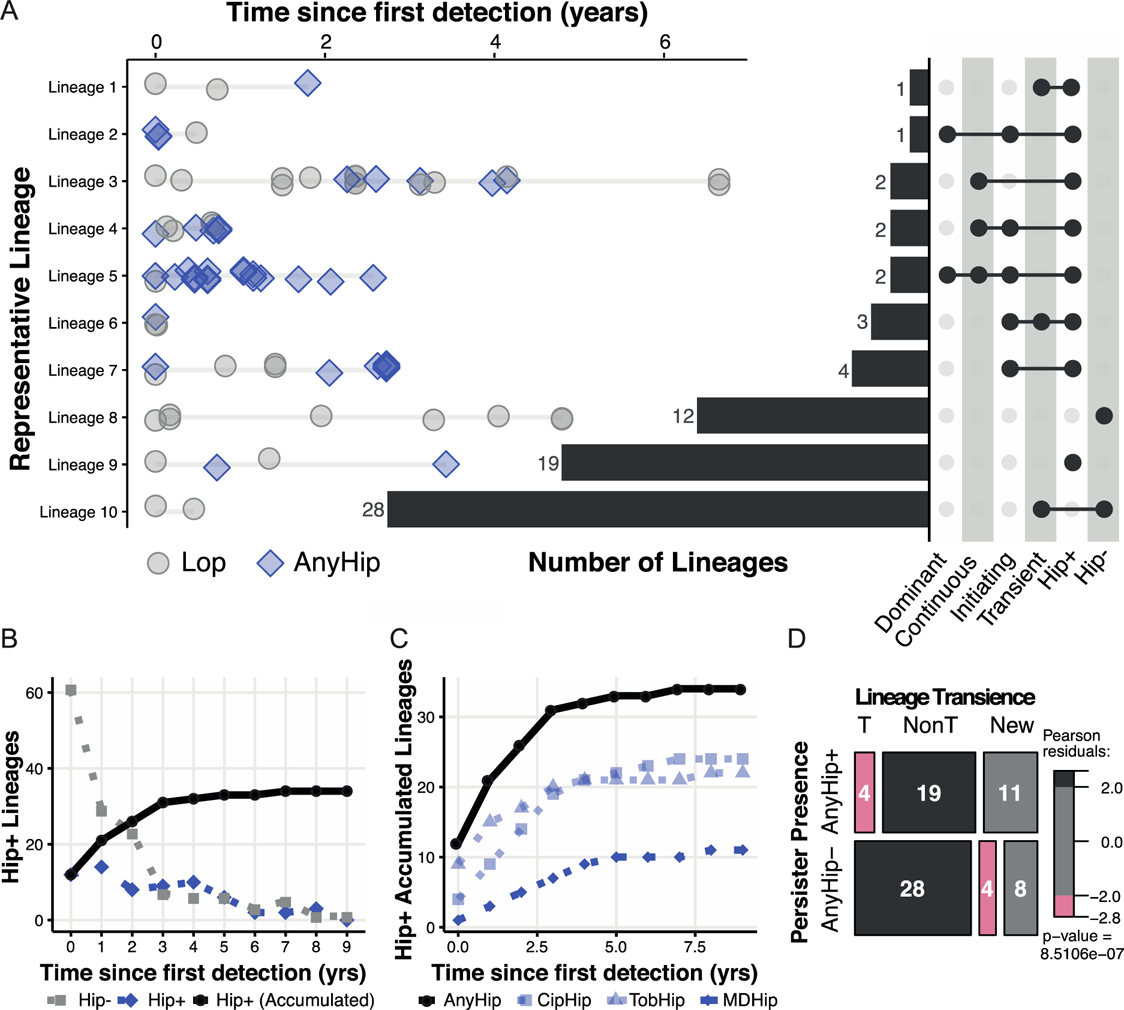
Hip Persister incidence patterns from a lineage-based perspective. **(A)** Lineages were classed according to several nested characteristics: transient versus non-transient lineages, Hip presence (AnyHip, meaning Cip Hip, Tob Hip or MD Hip), continuous periods of isolated Hips, lineage-initiating Hips, and Hips dominating a lineage. Lineages representing each combination of traits are shown on the left (Hip blue diamonds, Cip grey circles), while characteristic sets are identified and enumerated for the entire collection on the right. **(B)** For the AnyHip dataset, the continuous patient count of Hip-patients (grey circles) versus Hip+ patients (blue diamonds) for the prior year of colonization is plotted, while the accumulating count of Hip+ patients from time 0 is shown by black diamonds. **(C)** The accumulating count of Hip+ patients is shown for all four datasets (AnyHip as black circles, CipHip as transparent blue squares, TobHip as transparent blue triangles, and MDHip as dark blue diamonds). **(D)** Transient lineages (lineages of shorter than 2 years duration, less than 50% of total patient infection length, and which are followed by the appearance of a new lineage) are significantly associated with lineages lacking Hips (Hip-), while non-transient lineages are associated with the presence of Hips based on Pearson’s chi-squared test (via a mosaic plot visualizing a multi-way contingency table). Transience-unclassifiable lineages of shorter than 2 years’ duration at the end of a patient’s collection period are shown for context (‘New’).

### Hip variants accumulate over colonization time and occur rarely in transient lineages

We summarized the incidence of Any Hip variants over each lineage’s time of colonization in Figure 4B. Here we plot the continuous counts of patients exhibiting the Hip phenotype (Hip+) versus no Hip presence (Hip-) within the previous year (dashed lines) as well as the accumulation of lineages that have exhibited a Hip variant at least once by a certain age of colonization (solid line). The former illustrates both the number of lineages assessed at a given colonization age and the increasing fraction of Hip+ versus Hip-lineages over time. The latter shows how the likelihood of Hip emergence increases over time. Interestingly, when we plot Hip accumulation for all four datasets (Fig. 4C), we also observe twice as many lineages with initiating Hips in the Tob Hip dataset than the Cip Hip dataset, but Cip Hip incidence quickly catches up with and eventually exceeds Tob Hip incidence over time. Overall, Any Hip variants affect 34 lineages (Figure 4B-C) and 77% of the patients in our study cohort (Figure S2) by the end of the study period.

Next, we evaluated the relationship between lineage transience and Hip presence. We first mark the lineages present for less than 2 years in a patient at the end of their monitoring period as ‘New’ lineages since we cannot determine transience without additional samples. Of the 56 remaining lineages, non-transient lineages are significantly associated with the Hip+ lineage status, while transient lineages are significantly associated with the Hip-lineage status (Fig. 4D). Thus, a given patient often has multiple infecting lineages, but the Hip-lineages are much more likely to disappear over the course of infection. In summary, we find that despite variable incidence patterns, a clear majority of patients are infected by Hip+ lineages, and these lineages have a significant persistence advantage in comparison to Hip-lineages over time, suggesting that the Hip phenotype contributes to a fitness increase in antibiotic-treated patients.

### Hip variants show increased fitness in patient-similar biofilms

We next asked if Hip isolates are able to survive antibiotic treatment better than Lop isolates with similar antibiotic susceptibilities and growth properties in more complex conditions. We simulated antibiotic treatment of CF patients in a recently developed biofilm Pharmacokinetic/Pharmacodynamic (PK/PD) system, in which the bacteria are challenged with antibiotics in much the same way as in patients (38). We chose this model because *P. aeruginosa* often appears as biofilms in lungs of CF patients (39), because biofilms have been shown to harbor increased levels of persister cells (40), and because our model mimics the bacterial exposure to ciprofloxacin treatment as described for CF patients (38, 41). The isolates that were chosen shared similar 1) time since their first detection in the CF lungs were similar, 1. 2) MIC values for ciprofloxacin, 3) growth rates, and 4) belong to the same clone type. The Hip/Lop pair was differentially tagged with yellow fluorescent protein, YFP (Hip), or cyan fluorescent protein, CFP (Lop). Both strains formed biofilms with comparable biomasses in the flow-cell system. Hip and Lop cells were then mixed 1:1 and allowed to form a mixed biofilm. Representative images of the Hip/Lop biofilms are shown before and after treatment with ciprofloxacin (Fig. 5A).

**Figure 5.**
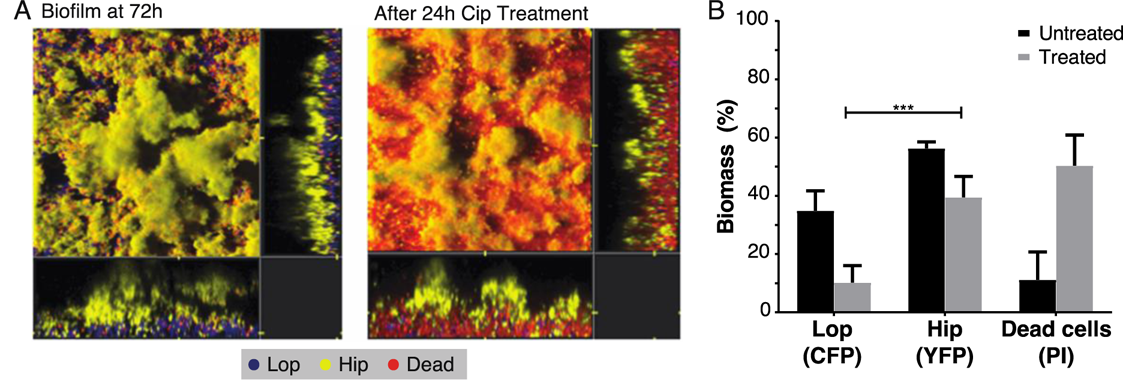
Fitness comparison of Lop and Hip isolates in biofilm conditions. **(A)** A representative Lop and Hip isolate with similar characteristics (Lop: cip MIC 1.0 µg/ml; growth rate 0.27 hr-1; time since first detection 4.28 years. Hip: cip MIC 0.75 µg/ml; growth rate 0.25 hr-1; time since first detection 5.49 years) were differentially tagged with CFP (Lop) or YFP (Hip). Tagged isolates were cocultured and allowed to form biofilms in a flow-cell model for 72 hours. Mixed biofilms were treated for 24 hours with ciprofloxacin (4 µg/ml). Propidium iodine (PI) was added to visualise dead cells (red). **(B)** Biomass was quantified for each population. Significant differences in biomass following treatment were determined using unpaired t-test (*** p <0.001).

The majority of Lop bacteria were located close to the glass substratum with the Hip population proliferating at the external surface of the biofilm, facing the liquid flow. The addition of ciprofloxacin preferentially killed the Lop population leaving the Hip population relatively unaffected by the antibiotic. COMSTAT analysis confirmed this changed population structure after ciprofloxacin addition (Fig. 5B). This documentation of a Hip associated fitness increase in an antibiotic containing environment is all the more striking, as it has been shown previously that ciprofloxacin treatment of flow-cell biofilms preferentially kills the surface sub-populations of micro-colonies (42) – yet the Hip cells on the colony surfaces survive much better than the internal Lop bacteria under treatment with ciprofloxacin.

## Discussion

We have mapped the prevalence of persisters in a large, aligned cohort of patients under intensive antibiotic treatment for a 10 year period (22, 41). Of 460 *P. aeruginosa* isolates from the airways of 39 young CF patients (74 lineages in total), 24.8% of the isolates were scored as robustly persisting Hip using a high-throughput screening approach to assay persistence against ciprofloxacin or tobramycin (Fig. 1). We show that the isolates display different levels of persisters, in accordance with the variance previously found between species and within strains (5, 43, 44). Most adaptive changes occur during the first few years of colonization (18, 45), which matches our objective of searching for signs of increased fitness of Hip variants in patients treated continuously with antibiotics. We show that in a young CF patient cohort impacted by early longitudinal colonization by *P. aeruginosa* strains, Hip variants were sampled from 77% of the patients (N=30) during a 10-year observation window. Our analysis is a new and important comparative baseline for developing effective surveillance, impact assessment, and eventual control of the persister phenotype in the clinic.

In the early years of infection after first detection of *P. aeruginosa* clone types, the Hip phenotype appeared and disappeared over time in our routine clinical sampling (Fig. 4A). While we see only partial overlap in Hip phenotype between our tobramycin and ciprofloxacin screens, the number of lineages and patients that exhibit Hip variants increases over time for all datasets (Fig. 4B-C, SI Appendix, Fig. S2), suggesting a selective advantage of this phenotype during the continuation of antibiotic therapy. In general, the majority of lineages that showed short-term colonization were made up of only Lop variants, which may partly explain why they were unable to establish a persistent infection (Fig. 4D). These in-patient data support the hypothesis that the Hip phenotype may generally have increased fitness in the antibiotic-containing lung environment. It is, however, important to note that neither dominance nor continuous presence of Hip variants is observed frequently (Fig. 4A). It is likely that fitness trade-offs and clonal interference impact on the fitness properties and the persistence level of Hip variants (14).

Multiple relationships between the Hip phenotype and other phenotypic traits such as growth rate and antibiotic resistance have been suggested in the literature. While some studies point out that there is no correlation between the mean growth rates of isolates and the observed Hip phenotype (46–48), reduced growth rates have been associated with high persister phenotypes in *E. coli* (28). A recent study in *Salmonella enterica* further supports that slow growth (regardless of mechanism) promotes the persister phenotype (31). We see that lineages which produce isolates with reduced growth rate are significantly more likely to also produce Hips. Furthermore, multiple genes targeted for mutation at a higher rate in Hip+ lineages are known to induce a growth defect (Table 1, SI Appendix, Dataset 2). However, we also observed the Hip phenotype among naïve, fast growing clinical isolates of *P. aeruginosa* (Fig. 3), supporting that other factors also influence the phenotype. Drug-tolerant cells have also been proposed to facilitate evolution of true antibiotic resistance in *E. coli in vitro* (49). Intermittent antibiotic exposure of a batch culture of *E. coli* selected for mutant clones harboring tolerance mutations that increased the growth lag-time, during which tolerance to killing by ampicillin selected for MIC-increasing mutations. Though *P. aeruginosa* in the CF lung is also exposed to fluctuating concentrations of antibiotic, our stringently defined Hip phenotype emerges simultaneously or after resistance in a majority of cases in contrast to these findings. We also observe more or less an equal number of lineages where Hip variants and resistant clones evolve independently in patients under antibiotic selection pressure, which has previously been suggested by comparative studies of lab strains (50). In summary, our results suggest that the Hip phenotype may be an early advantageous adaptation (18) arising stochastically in infected patients treated with antibiotics.

In many ways, investigations of the genetic underpinnings of persisters have been performed analogously to studies of antibiotic resistance, i.e. it was expected that a relatively limited set of genes defines the phenotype. In a study of urinary tract infection *E. coli* isolates, a gain-of-function mutation in the HipA toxin was commonly observed (21). In contrast, a lack of common targeted genes in a small collection of clinical Hip strains of *Mycobacterium tuberculosis* suggested utilization of multiple genetic pathways (51). Working at a much larger collection scale in a faster adapting organism, we do not see enrichment of mutations which previously have been associated with Hip phenotypes *in vitro*. A role for the top proposed persister target *aceF* has yet to be described, likely owing to its important role in growth; mutants with severe growth defects are often overlooked in genome-wide analyses *in vitro*. In support of this, an *aceF* transposon mutant is not available from the widely used PAO1 two-allele mutant library, and *aceF* did not appear in a recent persister screen of a pool of 100,000 unique PAO1 transposon mutants (20, 52). The *aceF* gene is mutated in 20% of the Hip+ lineages, which have been genotypically evaluated in our large, longitudinal collection, but in three of six lineages, it is also mutated in Lops. Meanwhile, no other enriched mutated genes are affected in as many Hip+ lineages alone (where only Hip isolates are affected). We therefore conclude that a Hip phenotype may derive from a diverse array of accumulating genetic changes, and it is likely that more than one mutation often determines the persister level in the respective bacterial populations. There are certainly many adaptive routes to slowed growth rate, which we have previously demonstrated is a convergent adaptive outcome in this early isolate collection (18). Our results likely reflect the multiple and dynamic selection pressures *in vivo*, which challenge Hip variants in antibiotic-treated populations very differently than those assessed in steady state *in vitro* conditions with only one selective force.

Many studies have shown the increased survival of persister cells under antibiotic treatment (28, 53) and then screened for genetic determinants of persistence, but few have evaluated the fitness of Hip versus Lop variants in direct competition experiments (43). In our study, we tested a Lop/Hip pair of isolates matched by genotype, phenotype, and colonization age in order to characterize the selective advantage of the Hip phenotype in a biofilm under treatment-replicating antibiotic exposure (38). We show that Hip cells survived ciprofloxacin treatment far better than Lop isolates and this survival is potentially reliant on biofilm architecture. Additionally, while homogeneous monoclonal *P. aeruginosa* biofilms treated with ciprofloxacin show preferential killing of bacteria in the top layers (14, 42, 54), Lop bacteria are preferentially killed in the deeper layers of the biofilm, showing an unexpected phenotype worthy of further study. It is also striking that the *in vitro* biofilm fitness assessment shows efficient elimination of the Lop strain in the presence of ciprofloxacin, whereas Hip variants often coexist with Lop variants *in vivo* (Fig. 4A). This suggests that in the patient, direct competition is likely limited by the large lung volume, many separate regional niches, and influence of the host (55).

In summary, we have shown that Hip variants of *P. aeruginosa* emerge frequently in young CF patients, and our results provide the first window into the evolving landscape of persistence across a whole patient cohort. As pathogens increase their fitness in patients over time, they clearly deploy the high persister phenotype as an important component in their survival repertoire and can do so from the earliest stages of infection. It is still premature to conclude that the high persister phenotype described here differs from what has been identified as Hip in *in vitro* experimental conditions, but we consistently find a much broader bacterial repertoire for survival in patient lungs. Hip variants do not seem to be mutated in genes previously found from *in vitro* experiments to associate with Hip or in any strongly conserved genetic route. We suggest that the difference in complexity of selection pressures when comparing *in vitro* and *in vivo* environmental conditions results in highly different evolutionary trajectories. With our investigation, we provide an important platform for broader clinically based studies and contribute important new context for monitoring and one day hopefully preventing the high persister phenotype in the clinic.

## Materials and Methods

### Strain collection

In total, we analyzed 460 *P. aeruginosa* airway isolates from young CF patients followed at the Copenhagen CF-clinic at Rigshospitalet (Dataset 1). The local ethics committee at the Capital Region of Denmark (Region Hovedstaden) approved the use of the stored *P. aeruginosa* isolates: registration number H-4-2015-FSP. Phenotyping data for 434 isolates of this strain collection (growth rate in LB, adhesion in LB, and ciprofloxacin MIC) have been previously published (18). We include additional tobramycin MIC measurements and expand the complete trait dataset to 446 isolates. All available trait data is provided in Dataset 1 along with the persister classification of each isolate and descriptive data.

Of the 460 isolates examined in the study, 403 isolates from 32 patients were described previously in Marvig et. al. (22) and the remaining isolates were taken from seven previously undescribed patients. The isolates were collected and stored at the Department of Clinical Microbiology at Rigshospitalet, Copenhagen, Denmark, between 2002 and 2014. Of the patients included in this study, 35.9% were diagnosed as chronically infected with *P. aeruginosa* by the end of the study period. We defined chronicity based on the Copenhagen CF Centre definition, whereby either *P. aeruginosa* has been detected in six consecutive monthly sputum samples or fewer consecutive sputum samples combined with observation of two or more *P. aeruginosa*-specific precipitating antibodies (41, 56). Intermittently colonized patients were defined as patients where at least one isolate of *P. aeruginosa* is detected, and normal levels of precipitating antibiotics against *P. aeruginosa* were observed.

### High-throughput screening for Hip mutants

To determine the frequency at which *P. aeruginosa* Hip mutants emerge in CF patients, we screened 460 isolates for the ‘persister’ phenotype against either ciprofloxacin or tobramycin. Stock 96-well microtiter plates containing 4 technical replicates of each isolate stored in glycerol (25 % v/v) were prepared and stored at −80°C. Using a 96-well spot replicator, bacteria were transferred from the stock plates into sterile 96-well microtiter plates containing 150 μl of Lysogeny Broth (LB) media. Plates were incubated statically for 48 hours at 37°C until the bacteria reached the stationary phase of growth. To determine the initial viability of bacteria in each well, the replicator was used to spot bacteria onto LB agar plates. Subsequently, 100 μg/ml of either ciprofloxacin or tobramycin was added to each well and the microtiter plates were incubated statically for a further 20-24 hours at 37°C. Serial dilutions were performed in 96 well microtiter plates containing 0.9 % NaCl using an automated fluid handling robot (Viaflo3844/ Integra Biosciences AG). Each dilution was spotted onto LB agar plates using the replicator and plates were incubated at 37°C for at least 24 hours. The growth of the bacteria was compared by counting colonies whenever possible and visually inspecting growth on the plates before and after antibiotic treatment. Experiments were performed in duplicate for each antibiotic.

### Persister assay validation

Time-kill experiments were performed for six isolates from the same lineage (3 Hip and 3 Lop). *P. aeruginosa* were inoculated in 3 ml of LB media in 14 ml culture tubes and incubated for 48 hours at 37°C with shaking at 250 rpm. Following incubation, each culture was serially diluted using sterile 0.9 % NaCl, plated onto LB agar and incubated at 37°C to determine the initial colony forming units (CFU). The remaining culture was treated with 100 μg/ml of ciprofloxacin and incubated at 37°C with shaking. Cultures were washed and diluted in sterile 0.9 % NaCl, then spot plated onto LB agar 6 and 24 hours after the addition of antibiotic. Plates were incubated for 24 hours at 37°C. Bacteria survival was measured by counting CFU per ml. For an additional 19 isolates, the same validation experiment was performed, however cultures were only plated out after 24 hours, which was within the ‘persister plateau’.

### Phenotype screening

The same frozen library of isolates used in the persister screening was also replicated for assay of minimum inhibitory concentrations (MICs), bacterial growth, and adhesion as described below and in Bartell et al (18). MICs for ciprofloxacin and tobramycin were determined using E-test methodology according to the manufacturer’s recommendations (Liofilchem®, Italy). To assay growth rate, bacteria were replicated from frozen plates into 96 well plates containing 150µL of LB medium, and incubated for 20 hours at 37**°**C with constant shaking. OD 630 nm measurements were taken every 20 minutes using a microplate reader (Holm & Halby, Copenhagen, Denmark/Synergy H1). Generation times (Td) were determined on the best-fit line of a minimum of 3 points during exponential growth of the bacterial isolate. Growth rates (hr_-1_) were calculated using the formula log (2)/ Td x 60 using semi-automated code described in Bartell et al (18). Adhesion was measured via attachment assays in 96-well plates using NUNC peg lids and 96 well plates with 150µl Luria broth medium. OD_600nm_ was measured after incubation for 20 hours at 37**°**C and subsequently, a “washing microtiter plate” with 180µl PBS was used to wash the peg lids and remove non-adhering cells. After transfer of the peg lids to a microtiter plate containing 160µl 0.01% crystal violet (CV), they were left to stain for 15 min. To remove unbound crystal violet, the lids were then washed again three times in three individual “washing microtiter plates” with 180µl PBS. Adhesion was measured by detaching adhering CV stained cells through washing the peg lids in a microtiter plate containing 180µl 99% ethanol. An ELISA reader was then used to measure the CV density at OD_590nm_. (Microtiter plates were bought at Fisher Scientific, NUNC Cat no. 167008, peg lids cat no. 445497).

### Pharmacokinetic/Pharmacodynamic (PK/PD) flow chamber biofilm model

For fitness experiments, we used a PK/PD biofilm model system combined with confocal laser-scanning microscopy. This system simulates the changing antibiotic concentrations in CF patients during intravenous dosing in addition to retaining a similar profile of antibiotic decay as the one taking place in CF patients (38). First, Hip and Lop isolates were differentially tagged with a yellow fluorescent protein (YFP) or cyan fluorescent protein (CFP) respectively (14). Flow chambers were inoculated with a 1:1 mixture of Hip and Lop bacteria (each isolate had an initial OD_600_ of 0.5). Bacteria were incubated for one hour at 30 °C, then nutrient flow was applied to each chamber (40x diluted LB at a rate of 20 ml/h using a Watson Marlow 205S peristaltic pump). Biofilms were allowed to form for 72 hours, at which point flow was stopped and medium containing ciprofloxacin was added. Peak ciprofloxacin concentrations were calculated to be 4 mg/L based on PK parameters generated from healthy patients and CF patients (57). The medium was pumped from the dilution flask through the antibiotic flask to the flow chambers at a constant rate calculated to mimic the elimination rate constant of the antibiotic for 24 hrs. A confocal laser-scanning microscope (Zeiss LSM 510) equipped with an argon/krypton laser and detectors was used to monitor YFP (excitation 514 nm, emission 530 nm), CFP (excitation 458 nm, emission 490 nm), and dead cells (propidium iodine, excitation 543 nm, emission 565 nm). Multichannel simulated fluorescent projections (SFPs) and sections through the biofilms were generated using Imaris software (Bitplane AG, Switzerland). The images were later analyzed using COMSTAT (58). The PK/PD biofilm experiments were performed using two independent Hip/Lop isolate pairs. Pairs were taken from the same patient at a similar time since first detection and had similar growth rates and ciprofloxacin MICs (Table S2). The data presented are from 2 biological experiments with 4 independent images taken from each experiment.

### Lineage-based genetic analysis

To generate a list of mutated genes associated with the Hip phenotype, we used previously generated whole-genome sequencing data and variant calling filtered to obtain nonsynonymous mutations that had accumulated within a lineage after the first isolate (22) to evaluate differential mutation patterns for Lop and Hip variants for 403 sequenced isolates. In this filtering process, we also removed mutations associated with any known ‘hypermutator’ isolates based on a mutation in *mutS* or *mutL* to avoid the influence of high random mutation in these isolates on the analysis. To identify genes that were mutated more than would have been expected by drift/random mutation while accounting for lineage-based mutation accumulation over time, we adapted a statistical analysis of the relative mutation enrichment by lineage. After separating Lop and Hip variants, we compared the mutated-gene lineage enrichment ratios for each group - the number of lineages with observed mutation(s) in a given gene divided by the number of lineages expected to have mutations in that gene according to random mutation. This enrichment metric was obtained as follows for each group: we determined the observed number of lineages mutated (sum-obs) in each gene. Then we estimated the average number of lineages (avg-exp) that would have been mutated in each gene if mutations were spread out randomly over the PAO1 genome. Using a random-roulette algorithm, the number of genes that were observed to be mutated in a given lineage was spread out over the PAO1 genome for 1000 iterations, providing a *m*_gene_ by *n*_iteration_ matrix of randomly mutated gene profiles for each lineage. For the same iteration *n* across all lineages, it was noted whether a given gene was mutated. This allowed us to determine an average number of lineages expected to be mutated over 1000 iterations. If a gene was hit by chance more than once in a single iteration, this would still only be denoted as one hit; this is in alignment with our observed mutation assessment, where multiple isolates could be hit in the same gene but we only noted whether or not the lineage was hit by unique mutations in the specific gene. After obtaining the relative enrichment by lineage, a Poisson distribution was used to calculate the probability of the observed given random drift (expected). We also divided the lineage enrichment metric for genes mutated in Hip variants by that for Lop variants to obtain a lineage enrichment ratio to identify targeted genes particularly impactful in the evolution of the Hip population.

### Data analysis and statistics

Analyses were conducted in RStudio v. 1.0.143 and R v. 3.4.0 with visualization package ggplot2 v. 3.0.0. Lineage set analysis was performed using UpSetR v. 1.3.3 in R (59). Principal component analysis was performed in R using ‘prcomp’ with centered and scaled phenotype data (Dataset 1). Mosaic plots (visualizing multi-way contingency tables) showing the association between two variables via the conditional relative frequency and significant associations based on a Pearson X_2_ test were created using vcd v. 1.4-4 in R (60).

### Data availability

Screen data for the persister phenotype is provided as a supplemental dataset (Dataset 1) including trait data, which partially overlaps with trait data published previously (18). Code in R for mosaic plots, PCA analysis, and set intersection analysis of lineage characteristics is available on request. Lineage enrichment analysis was performed in Anaconda3 v. 4.0.0 using custom scripts available on request.

## Acknowledgments

This work was supported by Cystic Fibrosis Foundation Pilot and Feasibility Award to KL. HKJ was supported by The Novo Nordisk Foundation (NNF12OC1015920 and NNF15OC0017444), by Rigshospitalets Rammebevilling 2015-17 (R88-A3537), by Lundbeckfonden (R167-2013-15229), by RegionH Rammebevilling (R144-A5287) and by Independent Research Fund Denmark (DFF-4183-00051). JAB was supported by postdoctoral fellowships from the Whitaker Foundation and the Cystic Fibrosis Foundation (BARTEL18F0). We thank Katja Bloksted, Ulla Rydahl Johansen, Helle Nordbjerg Andersen, Sarah Buhr Bendixen, Camilla Thranow, Pia Poss, Bonnie Horsted Erichsen and Rakel Schiøtt for excellent technical assistance at Rigshospitalet. We thank Prof. Vasili Hauryliuk for helpful comments.

## Author Contributions

SM and KL designed the study. HKJ collected all the bacterial isolates. BM, DRC, JAB, and JAJH performed all experiments. JAB and LMS performed genetic and lineage-based data analysis. All authors contributed to the writing of the manuscript. All authors approved the final version.

## Ethics approval

The local ethics committee at the Capital Region of Denmark (Region Hovedstaden) approved the use of the stored *P. aeruginosa* isolates: registration number H-4-2015-FSP. We confirm that all methods were performed in accordance with the relevant guidelines and regulations.

## Supplementary information

**Supplementary methods.** Description of phenotyping methods, biofilm flow chamber experiments, and lineage enrichment analysis.

### Dataset Descriptions

**Dataset 1. Isolate collection data, metadata, and screen results.** Description of all isolates including age, genotypic, and phenotypic information used in the analysis. Identification of Hips in CipHip, TobHip, MDHip, and AnyHip datasets as well as Persister Scores in each antibiotic screen. Specific legend included below and in dataset file.

ID: The id of the isolate.

Sequence_ID: Sequence id of isolate for internal use and can be referenced to the isolate collection published and analyzed in Marvig et al.: “Convergent evolution and adaptation of Pseudomonas aeruginosa within patients with cystic fibrosis”, Nature Genetics 47, (2015).

SRA_accNo: Accession number for sequences published previously in Marvig et al.: “Convergent evolution and adaptation of Pseudomonas aeruginosa within patients with cystic fibrosis”, Nature Genetics 47, (2015).

Genotype: The clone type/genotype of the isolate.

Patient: The patient wherefrom the isolate has been sampled.

Lineage: Patient-Genotype combination distinguishing specific clone types evolving in a given patient.

Date_no: Date of sampling, using fraction of year.

IageCT: The “Infection age of the Clone Type”, the time (in years) since the clone type of the specific isolate was first detected in the patient from which the specific isolate was sampled.

This is referred to in the study as “the time since first detection” or “colonization time” or “age of the lineage”.

Length1PA: The time (in years) since P. aeruginosa was first positively cultured from the patient’s lungs, regardless of clone type.

PersisterScore: Score of 0-4 according to minimum number of spotted cultures that grew after antibiotic exposure (minimum in 2 biological replicates of 4 spots, 8 total spots).

cip: MIC of Ciprofloxacin.

tob: MIC of Tobramycin.

AdhesionN: OD of crystal violet normalised against 20h of growth.

GR_LB: Growth rate in LB, h-1.

cipR: Isolate classification as ciprofloxacin sensitive or resistent based on EUCAST breakpoint of .5.

tobR: Isolate classification as tobramycin sensitive or resistent based on EUCAST breakpoint of 4.

hypermutator: designation of known hypermutator isolates based on mutations in mutL or mutS.

Persister: Hip versus Lop classification as 1 versus 0 using a cutoff of at least 3 spots growth in each biological replicate.

Persister_CIP: persister class from ciprofloxacin screen alone.

Persister_TOB: persister class from tobramycin screen alone.

Persister_MD: persister class from ciprofloxacin screen and tobramycin screen.

Hips must have scored as 3 or greater in both screens, representing a multidrug persister.

Persister_ANY: persister class from ciprofloxacin screen and tobramycin screen.

Hips must have scored as 3 or greater in at least one screen.

**Dataset 2. Lineage-based mutation enrichment analysis for coding genes and noncoding regions for each dataset (CipHip, TobHip, MDHip, AnyHip).** Mutated genes enriched in Hip versus Lop isolates as assessed from a convergent evolution perspective accounting for lineage adaptation. Lineage enrichment ratio was calculated by dividing lineage-based gene mutation enrichment within Hip variants by that within Lop variants for each gene. Top Hip-linked genes were selected via the following criteria: greater than 2 lineages presenting mutations in that gene in the Hip population and a lineage enrichment ratio greater than 2.

**Figure S1.**
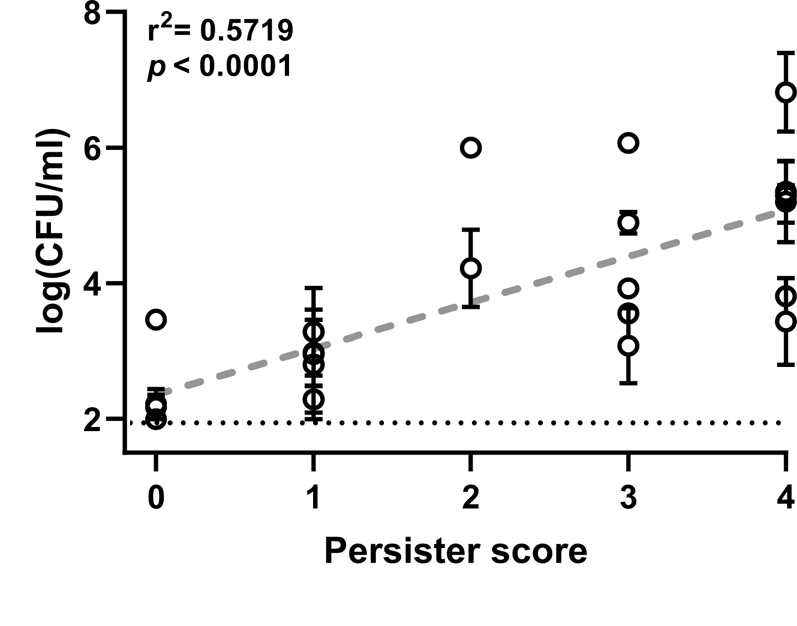
Correlation between persister score and colony forming units following treatment with ciprofloxacin. *P. aeruginosa* isolates representing each of the scores possible from the high-throughput screen (0-4) were treated with 100 µg/ml of ciprofloxacin for 24 hours, then plated on agar for surviving CFU determination. Each isolate was tested independently at least 4 times. The data are represented by the mean and SEM.

**Figure S2.**
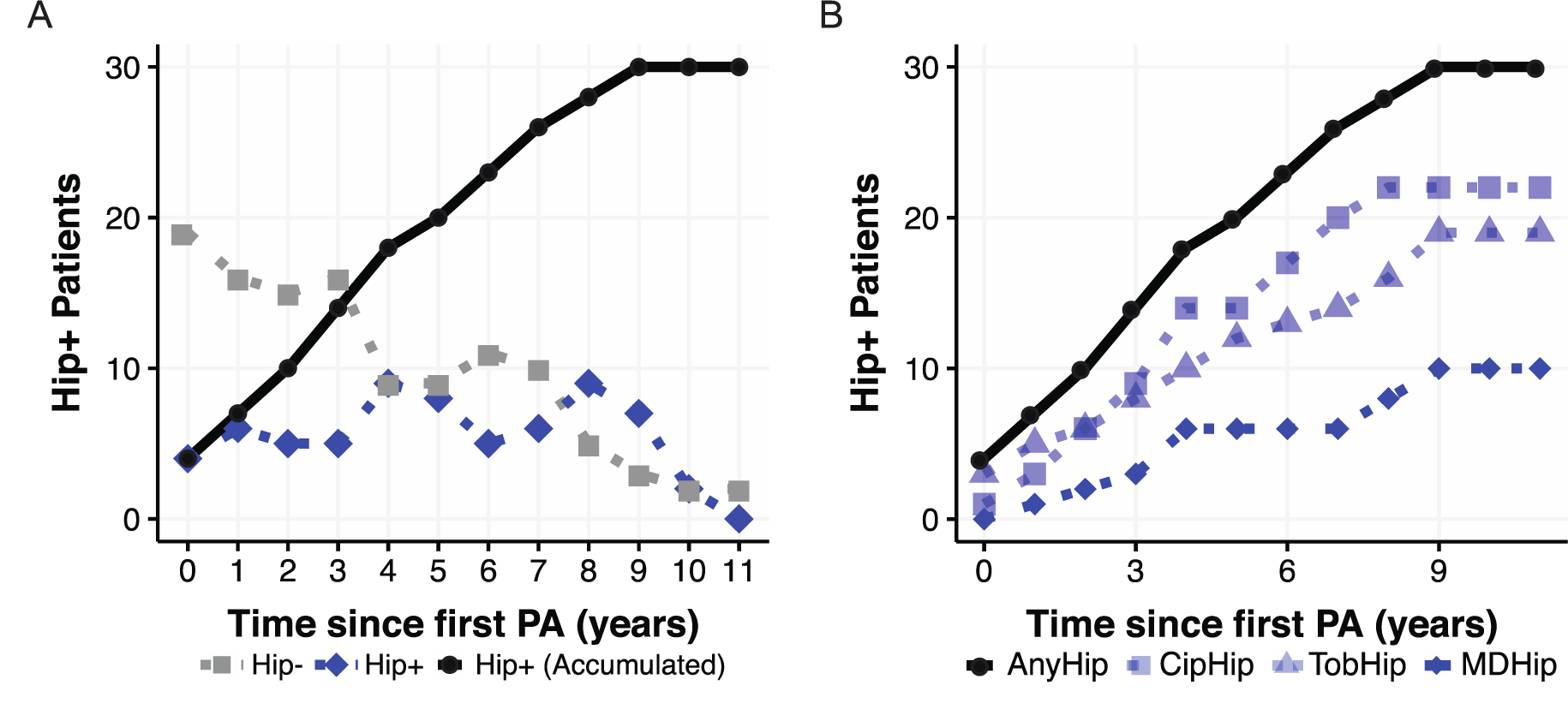
Hip accumulation in patients. (A) Hip-(gray squares) and Hip+ (blue diamonds) show the continuous count of patients with Hip-lineage(s) versus Hip+ lineage(s) for the prior year of colonization, while the accumulating count of patients with Hip+ lineages from time 0 is shown by black circles (AnyHip). **(B)** Accumulated patients with Hip+ lineages are shown for all persister datasets.

**Table S1.** Emergence of resistant versus persister isolates across 74 lineages via assessment of AnyHip isolates (persister score greater than 2) versus AnyHip plus transition isolates (persister score greater than 0).

| Persister<br>score | Category | # Lineages affected |  |  |
| --- | --- | --- | --- | --- |
|  |  | *Cip* | *Tob* | *Any* |
| **AnyHips**<br>**(Persister**<br>**score > 2)** | *Persisters alone* | 2 | 16 | 16 |
|  | *Resistance alone* | 14 | 4 | 14 |
|  | *Simultaneous emergence* | 11 | 1 | 11 |
|  | *Persisters before Resistance* | 2 | 3 | 5 |
|  | *Resistance before Persisters* | 8 | 2 | 9 |
| **AnyHips**<br>**plus**<br>**transition**<br>**variants**<br>**(Persister**<br>**score > 0)** | *Persisters alone* | 10 | 45 | 48 |
|  | *Resistance alone* | 8 | 1 | 8 |
|  | *Simultaneous emergence* | 7 | 2 | 7 |
|  | *Persisters before Resistance* | 15 | 7 | 15 |
|  | *Resistance before Persisters* | 5 | 0 | 5 |

**Table S2.** Persister genes identified in previous *P. aeruginosa* studies which we did not observe as targets in our lineage enrichment analysis.

| Locus | Gene | Product | Function | Reference |
| --- | --- | --- | --- | --- |
| PA5338 | *spoT* | Guanosine-3',5'-bis(diphosphate) 3'-phyrophosphohydrolase | Adaptation, protection; Nucleotide biosynthesis and metabolism | (8) |
| PA0934 | *relA* | GTP pyrophokinase | Adaptation, protection | (8) |
| PA3622 | *rpoS* | Sigma factor RpoS | Transcription | (9, 10) |
| PA4723 | *dksA* | DksA | Transcription; DNA replication | (8) |
| PA5002 | *dnpA* | de-N-acetylase | Membrane proteins | (11) |
| PA1045 |  | Hypothetical protein | Unknown | (11) |
| PA0299 | *spuC* | Polyamine:pyruvate transaminase | Carbon compound catabolism | (11) |
| PA3589 |  | Probable acyl-CoA thiolase | Carbon compound catabolism | (11) |
| PA3883 |  | Probable short-chain dehydrogenase | Putative enzyme | (11) |
| PA5261 | *algR* | AlgR | Two-component regulatory system; secreted factors | (11) |
| PA0318 |  | Hypothetical protein | Putative enzymes | (11) |
| PA0409 | *pilH* | PilH | Motility, chemotaxis; Two-component regulatory system | (11) |
| PA3166 | *pheA* | Chorismate mutase | Amino acid biosynthesis and metabolism | (11) |
| PA5332 | *crc* | Catabolite repression control protein | Carbon compound catabolism; energy metabolism | (12) |
| PA4756 | *carB* | Carbamoylphosphate synthase | Nucleotide metabolism; Amino acid biosynthesis and<br>metabolism | (13) |

